# The impact of mechanical requirements on the neural control of skeletal muscle and subsequent energetic rates

**DOI:** 10.64898/2025.12.04.692254

**Authors:** Ryan N. Konno, François Hug, Glen A Lichtwark, Taylor J.M. Dick

## Abstract

The energetic cost of skeletal muscle contraction is a fundamental driver in the selection of locomotor strategies. Muscle energy consumption depends on muscle-fibre typology, neural drive, and mechanical state. While the influence of fibre-type and mechanical state on energy use has been extensively characterised in isolated muscle preparations, these experiments fail to capture the influence of realistic *in vivo* motor unit recruitment strategies. Hence, this study aims to capture neural drive in the tibialis anterior, in particular motor unit recruitment thresholds and rate coding, and the corresponding changes in energy use across a range of mechanical demands. Fixed ankle angle dorsiflexion contractions were performed at varying rates of torque development, while high density electromyography characterized motor unit spiking activity, B-mode ultrasound captured muscle fascicle dynamics, and indirect calorimetry measured energetic rates. Faster rates of torque development required earlier recruitment of motor units with increased motor unit firing rates which coincided with increased fascicle strain rates. At higher rates of torque development, these altered motor unit discharge patterns and muscle mechanics coincided with increased muscle energy use. Together, these findings highlight that *in vivo* muscle energetics emerge from the dynamic interplay between neural control and mechanical demands.

## 2 Introduction

Skeletal muscle energy use is a prime determinant in the selection of locomotor strategies adopted by humans and other animals [1]. Existing literature has captured, in detail, the neuromechanical determinants of energy use during muscle contractions across a range of species and muscle types, but these experiments are typically performed in maximally activated isolated muscles or fibre-bundle preparations [2,3]. These conditions differ substantially from the submaximal muscle contractions that occur during everyday movements. *In vivo*, skeletal muscle force production is controlled by a pool of motor units, where each motor unit consists of a motor neuron and the muscle fibres it innervates. These motor units are typically recruited following an orderly fashion with the smallest motor neurons (low threshold) recruited before the larger motor neurons (high threshold) [4]; however, external stimulation results in an unnatural or possibly reversed recruitment order [5]. Thus, much of our knowledge regarding muscle energy use comes from experiments that do not reflect natural neural control. To ensure accurate extrapolation of energetic theories to *in vivo* muscle contractions, it is necessary to quantify muscle energetics in muscles with mixed fibre-type proportions, at submaximal activation that follow natural motor unit recruitment.

Skeletal muscle energy use, characterised as the energy liberated from a muscle contraction into mechanical work or heat, is the summation of three main energy consuming processes: cross-bridge cycling, Ca^+2^ transport costs, and regeneration of adenosine triphosphate (ATP) [3]. The energetic efficiency of these processes depends on the muscle-fibre type, as fibres with slow-type myosin isoforms are twice as efficient as faster muscle fibre types [6]. Given that muscle fibre-types tend be distributed such that lower threshold units correspond to slow-type fibres and higher threshold units to fast-type fibres [7], there is likely a role of recruitment strategy on the energetics of muscle contraction [8], but this remains largely unexplored.

It is well established that muscle energy use increases with higher levels of neural drive or electrical stimulation [2], but this can be amplified by changes in a muscle’s mechanical state. Muscle mechanical state (i.e., strain and strain rate) can influence the level of neural drive it receives; for example, producing a given muscle force at faster shortening velocities requires higher levels of neural drive [9]. Beyond influencing neural drive, mechanical state alters the efficiency of energy consuming processes [3]. Isolated fibre-bundle preparations have shown more energy is consumed at faster muscle shortening rates, independent of changes to neural drive [10]. The combined influence of muscle strain and strain rate has been summarised in existing phenomenological models of muscle energy use (e.g. [8,11]). While these relationships were first derived from isolated muscle preparations, human experiments have also shown that the dependence of energy use on mechanical state persists *in vivo* [12–15]. Despite this, energetic models still fail to accurately predict energetic rates during *in vivo* contractions [15,16]. One potential reason is that most models lack a detailed representation of the control of motor units [17], in part because experimental datasets that combine motor unit activity with simultaneous measures of muscle energetic rates do not yet exist.

This study aimed to simultaneously measure changes in *in vivo* motor unit activity and overall energetic rate across different rates of torque development, in order to establish a mechanistic link between muscle activation requirements, mechanical behaviour, and energetics. To quantify motor unit discharge characteristics (recruitment thresholds and firing rates), high density surface electromyography signals were decomposed to identify the spiking activity of a large number of motor units; providing a more representative estimate of the neural drive than traditional intramuscular recordings which sample a limited number of motor units [18]. Muscle fascicle dynamics were measured using B-mode ultrasound, while energetic rates were captured using indirect calorimetry. We hypothesised that higher rates of torque development would be associated with greater muscle fascicle strain rates, and, owing force-velocity constraints, a higher neural drive to the muscle. This higher neural drive would be identified by decreases in motor unit recruitment thresholds and increases in firing rates of matched motor units, and accompanied by increases in energetic rates. The outcomes are ultimately important for elucidating patterns of *in vivo* neural control and their contributions to energetic rates.

## 3 Methods

Data was collected in 18 participants (7 female, 11 male, 26.6±9.5 years, 177±9.6 *cm*, and 75.4±12.8 *kg* (Mean±SD)). Each participant provided written informed consent. Ethics was approved by the University of Queensland’s Human Research Ethics Committee (2022/HE001089). Data from one participant (P04) was removed due to the inability to extract motor units and energetic rates.

### 3.1 Protocol

Participants were instructed to fast for 12 hours prior to the experiment. The setup consisted of participants laying supine with their foot strapped into a custom-made ankle dynamometer with a flexed knee (mean knee angle: 135±6.5°, 180° being the knee extended) and slightly plantarflexed ankle (mean ankle angle: 97±6.8°, 90° being the angle between the tibia and the torque plate). Preliminary experiments revealed this position reduced coactivation of other lower limb muscles during dorsiflexion contractions (Figure 1). Dorsiflexion contractions limited the need to consider force-sharing strategies between muscles as the tibialis anterior (TA) is largely responsible for ankle dorsiflexion torque [19,20]. Following warm up contractions, participants performed three maximum voluntary contractions (MVC) with one minute rest between each contraction, with the highest intensity contraction used to normalise both torque and electromyography (EMG) data.

**Figure 1:**
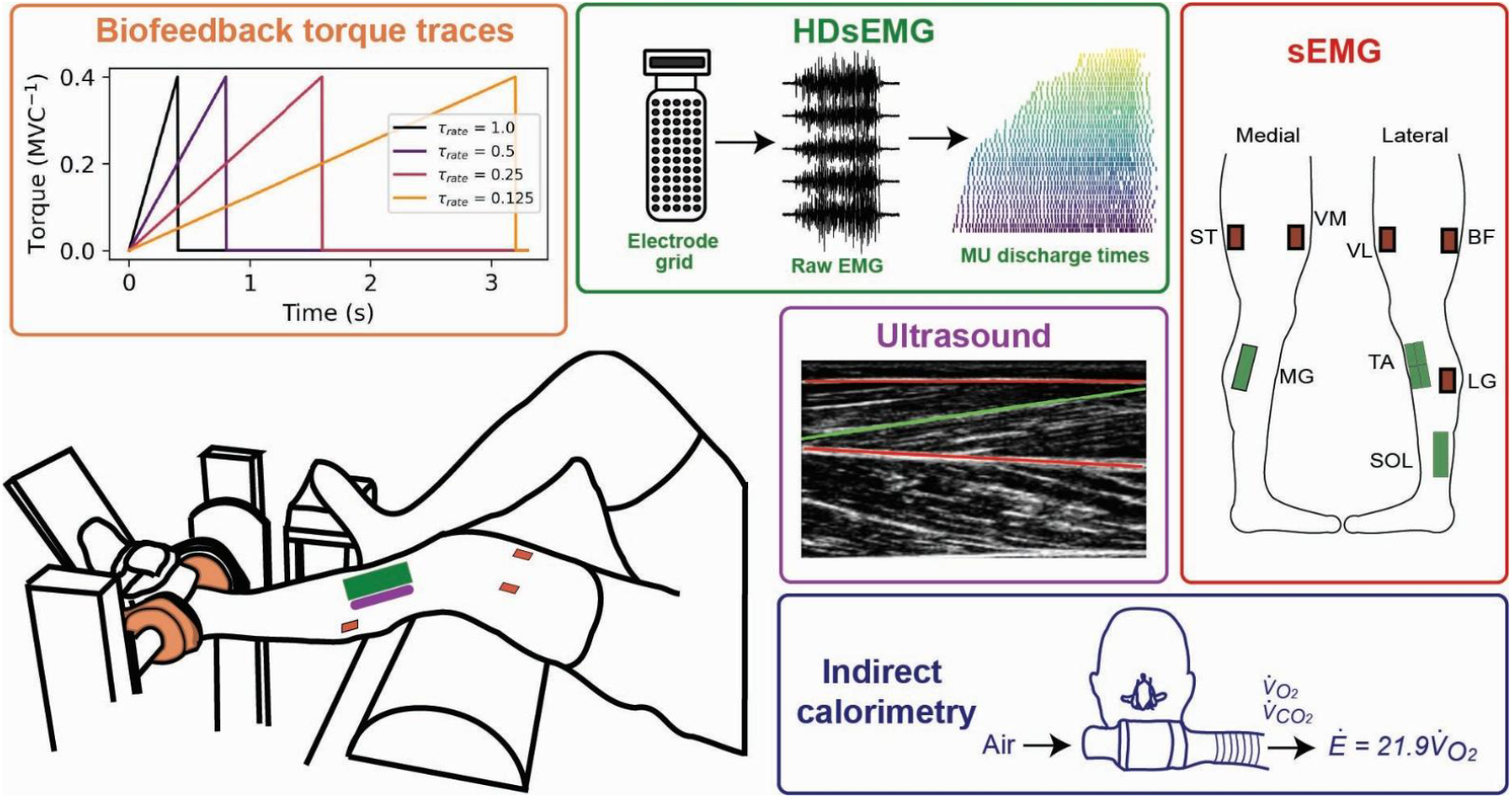
Experimental setup and analysis. Participants were positioned supine with their knee slightly flexed and ankle slightly extended and performed fixed ankle angle dorsiflexion contractions at varying rates of torque development 0.125, 0.25, 0.5, and 1.0 τ_mvc_ s^−1^. For each condition, contractions were performed for 5 minutes with rest periods of 1.5 × the time to reach 40% MVC to maintain constant torque integral across conditions. High density electromyography (HDsEMG) placed on the tibialis anterior (TA), medial gastronemius (MG), and soleus (SOL) measured TA motor unit discharge characteristics. Energetic rates were simultaneously measured using indirect calorimetry, while surface electromyography (sEMG) on the vastus lateralis (VL), vastus medialis (VM), semitendinosis (ST), biceps femoris (BF), and lateral gastronemius (LG) was used to measure co-activation. For the same conditions, B-mode ultrasound was used to quantify TA fascicle dynamics.

We simultaneously measured high-density electromyography (HDsEMG) in the TA, while indirect calorimetry measured whole-body energetic rate (Figure 1). The protocol consisted of four trials with prescribed ankle torque biofeedback provided visually via a monitor, with ramping contractions to 40% of MVC torque, *τ*_*mvc*_, at rates of torque development of 0.125, 0.25, 0.5, and 1 *τ*_*mvc*_ *s*^−1^. Contractions in each trial were performed repeatedly for 5 minutes to obtain stable energetic rates with the torque trace chosen to maintain a constant torque integral across each trial (Figure 1). Rest periods between consecutive contractions were 1.5× the time to reach 40% of *τ*_*mvc*_, which corresponds to relaxation times of 4.8, 2.4, 1.2, and 0.6 *s* for the 0.125, 0.25, 0.5, and 1 *τ*_*mvc*_ *s*^−1^conditions, respectively. Trials were performed in a random order with a minimum of 5 minutes of rest between trials to ensure a return to resting energetic rates before the next trial. Prior to the first trial, participants were given a familiarisation period at each rate of torque development to ensure they could follow the biofeedback torque trace accurately. B-mode ultrasound of the TA was measured under the same torque feedback conditions, but in separate 1-minute trials owing to space constraints on the TA as the EMG electrode grids covered most of the muscle surface.

### 3.2 Data Collection and Analysis

#### 3.2.1 High density electromyography

Four HDsEMG grids of 64 electrodes (4mm inter-electrode distance, GR08MM1305, OT Bioelettronica, Italy) were placed on the TA over the largest portion of the muscle belly with the length of the electrodes aligned parallel to the muscle shortening direction. The grids were positioned in two rows of two grids. This configuration was chosen to maximise the number of decomposed motor units [21]. One grid of 64 electrodes (8mm inter-electrode distance, GR08MM1305, OT Bioelettronica, Italy) was also placed on both the soleus (SOL) and medial gastronemius (MG). Prior to application of the electrode grids, the skin was shaved and cleaned using abrasive gel (Nuprep, Weaver and company, USA). The grids were attached to the skin using adhesive foam pads (OT Bioelettronica, Italy) with conductive paste (AC cream, Spes Medica, Genova, Italy) to ensure contact between the skin and the electrodes. A strap ground electrode was placed on the contralateral ankle, while a reference electrode (5 × 5 cm, Kendall Medi-TraceTM, Canada) was placed on the tibia of the tested leg. The EMG signals were recorded at 2048 *Hz* in monopolar mode and bandpass filtered at (20-500 *Hz*). The data were digitised and synchronised to the torque signal using the OT Bioelettronica Quattrocento system (EMG-Quattrocento; 400-channel EMG amplifier, OT Bioelettronica, Italy).

The HDsEMG signals were decomposed using a fast independent component analysis as implemented in MUedit [22]. Any poor-quality channels identified via visual inspection were removed from the analysis before automatic decomposition. The automatic decomposition was performed independently for each grid on the TA on a subset of the contractions over the 3-to 4-minute period of the trial. This subset was selected based on the contractions showing the highest *r*^2^ values between the produced torque and the ideal torque trace during the ramping phase of the contraction. The number of contractions included was the minimum required for reliable decomposition, while retaining only contractions with a high *r*^2^. Consequently, automatic decomposition was performed on fewer contractions for conditions at lower rates of torque development. This resulted in 4 contractions for the 0.125 *τ*_*mvc*_ *s*^−1^condition, 6 contractions for 0.25 *τ*_*mvc*_ *s*^−1^condition, and 8 for the 0.5 and 1.0 *τ*_*mvc*_ *s*^−1^conditions. After automatic decomposition, motor unit spike trains were manually edited following existing guidelines [23]. Identified motor units were tracked across trials by applying motor unit filters obtained after manual editing to the same grid in the other trials [21]. For example, motor unit filters obtained from trial A (e.g., 0.125 *τ*_*mvc*_ *s*^−1^) were applied to trial B (e.g., 0.25 *τ*_*mvc*_ *s*^−1^). The new motor unit discharge times obtained for trial B with the filter from trial A were then compared with the discharge times from the motor units obtained via direct automatic decomposition to trial B. A motor unit was successfully tracked if 30% of the discharge times were in common [21]. All motor units were included in the analysis regardless of whether they were tracked across all conditions, and any duplicate motor units across grids were removed.

Motor unit recruitment thresholds were calculated as ankle torque at the first motor unit discharge time. Values were averaged over the three contractions with the highest *r*^2^ between the produced torque and the ideal rate of torque development. The torque signal was lowpass filtered at 6 *Hz* and normalised to the MVC torque value. Motor units were categorised as either lower, medium, or higher threshold, based on the normalised recruitment threshold in the slowest contraction rate condition where the motor unit was detected. The corresponding bins for lower, medium, and higher threshold categories of [0, 0.13*τ*_*mvc*_] [0.13*τ*_*mvc*_, 0.27*τ*_*mvc*_], and [0.27*τ*_*mvc*_, 0.4*τ*_*mvc*_], respectively. Of note, these threshold categories are arbitrary and should be interpreted relative to the submaximal torque studied, where not all motor units were recruited. It is therefore unlikely that the highest threshold units were active, which was intentional in our design to limit the contribution of anaerobic pathways.

The motor unit firing rates were smoothed using a 400 *ms* Hanning window applied to the instantaneous firing rate [24]. From the smoothed trace, mean and peak firing rates were calculated over the cycle from the same contractions as those used to calculate the recruitment thresholds. To assess the relationship between motor unit firing rates and the rate of torque development over the contraction cycle, mean motor unit firing rates were computed by taking the mean of the smoothed firing rates over phases of the ramp contraction from torques of 0.05 to 0.1 *τ*_*mvc*_, 0.1 to 0.15 *τ*_*mvc*_, 0.15 to 0.2 *τ*_*mvc*_, and 0.2 to 0.25 *τ*_*mvc*_. A motor unit was only included in the mean if the firing rates were above 5 *Hz*, approximately the minimum firing rate for TA motor units [21], for the duration of the torque range, i.e. the motor unit is active for the duration of the window. Motor unit firing rates at recruitment were calculated as the mean of instantaneous firing rate over the first four discharge times.

We estimated the overall muscle activation level by determining the amplitude of the interference EMG signals. To do this, we differentiated the monopolar HDsEMG signal and then converted to a global EMG value using a differential between two cross-shaped patterns on the grid spaced apart by one electrode (giving a distance between the centres of the cross pattern of 16*mm* for the grid on the TA and 32*mm* on the SOL and MG). For each electrode in the pattern, any poor-quality channels were removed and interpolated using the surrounding channels. The raw signal was band-pass filtered using a 4th-order Butterworth filter from 10 to 500 *Hz*. The mean signal was calculated for each cross-pattern and a differential taken between the mean signals. The differential signal was then rectified and low-pass filtered at 12 *Hz*. To obtain maximal EMG amplitude values from the HDsEMG signal, a moving mean of 500*ms* was taken over the unfiltered rectified signal from the MVC trial with the MVC value taken as the maximum of this signal. Mean EMG amplitude values were computed over the whole cycle period including both the contraction and the rest phase. Due to poor quality signal, data from SOL of P07, P09, P16 and MG of P16 was removed from global EMG analysis.

#### 3.2.2 Assessment of co-activation

To monitor activation in other lower limb muscles, surface electromyography (sEMG) was recorded (Figure 1) in the lateral gastronemius (LG), vastus lateralis (VL), vastus medialis (VM), semitendinosis (ST), and biceps femoris (BF) (10mm interelectrode distance; Trigno Delsys Inc., Natick, USA). The sEMG signals were collected at 2048 *Hz* using Spike2 (V10, CED Ltd, Cambridge, UK) and synchronised to ankle torque with a data acquisition board (CED 1401, Cambridge Electronic Design Ltd., Cambridge, UK). To limit co-activation that could contribute to the whole-body energetic rate, the raw sEMG signals were monitored and verbal feedback was given to minimise co-activation if increases in sEMG magnitude were observed.

To obtain MVC values to normalise the sEMG signals, a knee extension (for VL and VM) and knee flexion contraction (for ST and BF) were performed while the participant was seated with resistance to knee torque was applied manually. MVC contractions for the plantar flexors (LG, MG, SOL) was performed while laying supine in the same position as the protocol. Two MVC contractions were performed for each muscle group with 1 minute rest between contractions. EMG signals from the trial and the MVC contractions were post-processed in Matlab (R2021b, MathWorks Inc., Natick, USA) with a band-pass filter from 10 to 500 *Hz*, then rectified and low-pass filtered at 12 *Hz*. After filtering, the trial sEMG signals were normalised to the maximum of the processed MVC sEMG signals. Due poor quality of the EMG recordings, VL and VM of P08, and BF of P05 and P06 were removed from the sEMG analysis.

#### 3.2.3 B-mode ultrasound

B-mode ultrasound (ArtUs EXT-1H, REV: C1S, Telemed, Vilnius, Lithuania) with a linear probe (LV8-5L60N-2, 60 mm, 5–8 MHz, Telemed, Vilnius, Lithuania) was used to measure TA muscle fascicle dynamics. A custom-made external trigger was used to sync with the torque trace and maintain a fixed 120 *Hz* sample rate. Individual fibre tracking was performed using the UltraTimTrack software with manual editing [25]. Two contractions were selected from 30 seconds of repeated contractions based on the highest *r*^2^ values between the ideal ramp and the measured torque trace. The fascicle strain was calculated as

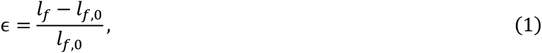

where *l*_*f*_ is the length of the fascicle and *l*_*f*,0_ is the resting length of the fascicle. The resting length was chosen independently for each contraction. The time varying fascicle lengths were computed for each contraction and lowpass filtered at 6*Hz*. Fascicle strain rates were calculated as the time derivative of the fascicle length data. Peak fascicle strain rates were defined as the maximum of the strain-rate over the torque ramp period. All fascicle measurements were taken from the superficial compartment of the TA. Participants P01, P02, P06, and P18 were removed from the ultrasound analysis due to an inability to identify either aponeurosis or fascicles.

#### 3.2.4 Indirect calorimetry

Whole-body energetic rates were obtained using indirect calorimetry (Vacumed Vista-MX2, Vacumetrics Inc., Ventura, USA) that was purposefully adapted with a low-flow rate setup (Flow sensor V70402, Vacumetrics Inc., Ventura, USA) to ensure steady measures near resting energetic rates. Standard indirect calorimetry setups for high flow rate exercise were not sensitive to the conditions in this study. Energetic rate was calculated using the mean 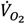 values over the 3-to 5-minute period of each contraction condition to ensure stable energetic rates. To convert from 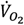 to energetic rate, we followed recommendations from [26], and used only 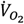 values, as whole-body 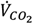 measurements are not representative of the muscle level metabolism. Thus, we calculated energetic rate as

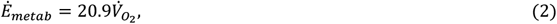

where 20.9 (*W mL*^−1^) gives the conversion between 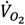 rate and energetic rate [27]. Rest periods were calculated prior to each trial with energetic rates averaged over 2 minutes prior to the trial once the energetic rates had stabilised (between 3 to 10 minutes of resting state). The resting energetic rate was then taken as the minimum of these resting rates. The net energetic rate for the trial was computed as Ė_*net*_ = Ė_*metab*_− Ė_*rest*_ and normalised to the participant body mass. Due to poor data quality, P02 and P17 were removed from the energetic analysis. The following trials were excluded because their average respiratory exchange ratio (RER) exceeded 1 over the 2-minute period: P08 and P15 at 0.125 *τ*_*mvc*_ *s*^−1^, P13 at 0.25 *τ*_*mvc*_ *s*^−1^, P08 at 0.5 *τ*_*mvc*_ *s*^−1^, and P12 at 1.0 *τ*_*mvc*_ *s*^−1^.

#### 3.2.5 Statistical analysis

All statistical analyses were performed in R (R Core Team, 2012) using lme4 (Bates, Maechler & Bolker, 2012) to run linear mixed-effects models to assess the relationship between rate of torque development and performed rate of torque development, mean global EMG, motor unit recruitment threshold, motor unit firing rate, peak fascicle strain and strain rates, and energetic rate. Rate of torque development was a fixed effect, while participant was a random intercept. Post hoc pair-wise comparisons were done with a Tukey correction as implemented in emmeans [28]. All results are reported as means, calculated as estimated marginal means using emmeans [28], with 95% confidence intervals. *r*^2^ values for the linear mixed-effects models represent conditional *r*^2^ values and *β* represents the fixed effect coefficient with respect to rate of torque development.

## 4 Results

### 4.1 Torque response

Participants were able to follow the torque bio-feedback across conditions (Table 1). The fastest condition (1.0 *τ*_*mvc*_ *s*^−1^) was the most difficult for participants to control, resulting in the largest deviation from the ideal rate of torque development (Table 1). No significant effect of condition was found on peak torque (*p* = 0.846). While the ideal torque trace was designed to maintain a constant torque integral across conditions, there was an effect of rate of torque development on the performed torque integral (*p <* 0.001, Table 1), which increased as rate of torque development increased (post-hoc *p*-values provided in supplementary material Table S1).

**Table 1:** Mean torque characteristics, fascicle dynamics, and energetic rates across participants.

| $\dot{\tau}_{ideal} (\tau_{mvc} s^{-1})$ | 0.125 | 0.25 | 0.5 | 1.0 |
| --- | --- | --- | --- | --- |
| $\dot{\tau}_{exp} (\tau_{mvc} s^{-1})$ | 0.13 [0.06, 0.20] | 0.27 [0.20, 0.34]* | 0.58 [0.51, 0.65]*# | 1.26 [1.19, 1.33]*#† |
| $\int \tau_{exp} (\tau_{mvc} s)$ | 29.8 [27.2, 32.4] | 33.5 [30.8, 36.1]* | 39.5 [36.9, 42.2]*# | 47.4 [44.7, 50.0]*#† |
| $\hat{a}_{max} (\% MVC)$ | 49.1 [41.4, 56.7] | 48.4 [40.8, 56.1] | 50.1 [42.5, 57.8] | 63.0 [55.3, 70.6]*#† |
| $\epsilon_{f,max} (unitless)$ | 0.14 [0.11, 0.17] | 0.13 [0.11, 0.16] | 0.13 [0.11, 0.16] | 0.13 [0.10, 0.16] |
| $\dot{\epsilon}_{f,max} (s^{-1})$ | 0.31 [0.24, 0.37] | 0.31 [0.25, 0.37] | 0.41 [0.32, 0.50] | 0.63 [0.46, 0.79]*#† |
| $\dot{E}_{net} (W kg^{-1})$ | 0.14 [0.02, 0.26] | 0.21 [0.09, 0.32] | 0.35 [0.24, 0.47]* | 0.44 [0.32, 0.55]*# |

$\dot{\tau}_{ideal}$ is the ideal torque rate and $\dot{\tau}_{exp}$ is the experimentally measured torque rate. $\int \tau_{exp}$ is the integral of the torque computed over the entire trial. $\hat{a}_{max}$ is the maximum activation over the trial. $\epsilon_{f,max}$ and $\dot{\epsilon}_{f,max}$ are the maximum fascicle strain and absolute value of the strain-rate, respectively. $\dot{E}_{net}$ is the mean energetic rate normalised to body mass. All values are reported as means with 95% CIs. \* denotes significantly different ( $p < 0.05$ ) than $0.125 \tau_{mvc} s^{-1}$ , # denotes significantly different ( $p < 0.05$ ) than $0.25 \tau_{mvc} s^{-1}$ , and † denotes significantly different ( $p < 0.05$ ) than $0.5 \tau_{mvc} s^{-1}$ .

### 4.2 Global EMG

Mean TA EMG amplitude increased with increases in rate of torque development (*β* = 0.095, *r*^2^ = 0.77, *p <* 0.001). Differences in maximum TA EMG amplitude were observed between the 1.0 *τ*_*mvc*_ *s*^−1^ condition and the 0.125, 0.25, and 0.5 *τ*_*mvc*_ *s*^−1^ conditions (all with *p <* 0.001, Table 1). There were small, but significant, increases in mean EMG amplitude of other lower limb muscles with increasing rates of torque development (*p <* 0.001 for VM, VL, ST, GM, GL, and SOL, and *p* = 0.012 for BF). Each muscle exhibited its highest mean EMG amplitude in the 1.0 *τ*_*mvc*_ *s*^−1^ condition; for the VM (*p* = 0.007), VL (*p* = 0.006), ST (*p* = 0.001), LG (*p* = 0.012), MG (*p <* 0.001), and SOL (*p <* 0.001), this value was significantly greater than in the 0.125 *τ*_*mvc*_ *s*^−1^condition. Across all conditions, the highest mean EMG amplitude was less than 2% of MVC EMG amplitude in the VM, VL, BF, and ST, and less that 5% in the and LG. SOL had the highest peak EMG amplitude of 5.8 [4.54, 7.07]% (see Supplementary Figure S1).

### 4.3 Motor unit discharge characteristics

A total of 263 unique motor units were detected across the 17 participants included in the analysis. Motor unit recruitment thresholds decreased with increasing rate of torque development for lower (*β* = −0.01 [−0.011, −0.004], *r*^2^ = 0.21, *p <* 0.001), medium (*β* = −0.037 [−0.043, 0.031], *r*^2^ = 0.40, *p <* 0.001), and higher (*β* = −0.06 [−0.072, 0.053], *r*^2^ = 0.57, *p <* 0.001) threshold motor units (Figure 2 B). There was a significant interaction between motor unit category (lower, medium, higher) and contraction condition on recruitment threshold (all *p* ≤ 0.029). Pairwise comparisons showed that recruitment thresholds decreased by 0.02 [0.003, 0.04] *τ*_*max*_ (*p* = 0.016), 0.12 [0.10, 0.14] *τ*_*max*_ (*p <* 0.001), and 0.20 [0.17, 0.23] *τ*_*max*_ (*p <* 0.001) from the 0.125 *τ*_*mvc*_ *s*^−1^ to the 1.0 *τ*_*mvc*_ *s*^−1^ conditions for the lower, medium, and higher threshold categories, respectively.

**Figure 2:**
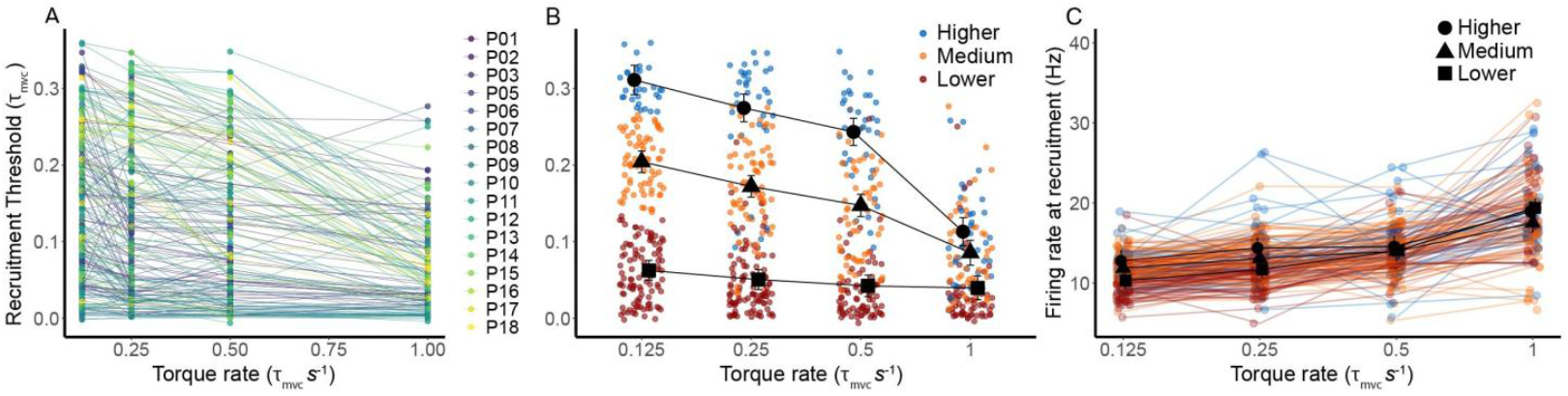
Motor unit recruitment threshold and firing rate varied across mechanical conditions. Motor unit recruitment threshold as a function of ideal torque rate, with each dot representing a motor unit and colours indicating individual participants (A). Recruitment threshold by motor unit category as a function of ideal torque rate, with each dot representing a motor unit and colours indicating individual motor unit categories (lower: 0 to 0.13 τ_mvc_, medium: 0.13 to 0.27 τ_mvc_, higher: 0.27 to 0.4 τ_mvc_) (B). Motor unit firing rate increased at time of recruitment as a function of rate of torque development (p < 0.001; C). Individual data points are the average of the firing rate between the first four motor unit discharge times. Lines represent tracked motor units in A-C. Means were calculated as estimated marginal means with 95% confidence intervals.

Firing rates at recruitment increased with rate of torque development, with a significant interaction between recruitment threshold category and rate of torque development (*p <* 0.001). The effect of rate of torque development on firing rate at recruitment was greatest in lower threshold units (*β* = 2.79 [2.53, 3.04]) compared to higher (*β* = 1.82 [1.25, 2.39]) or medium (*β* = 1.64 [1.28, 2.01]) threshold units (Figure 2C).

Mean motor unit firing rates increased with rate of torque development (*p <* 0.001). The relationship between motor unit firing rate and torque level is shown in Figure 3A-D. At lower torque levels, mean motor unit firing rates significantly increased with rate of torque development (*p <* 0.001; Figure 3E-H). Larger fixed effects of rate of torque development on mean firing rates were found at lower torque levels (0.05 to 0.1 *τ*_*mvc*_: *β* = 4.49 [3.44, 5.57]; 0.1 to 0.15 *τ*_*mvc*_: *β* = 5.11 [4.18, 6.04]; 0.15 to 0.2 *τ*_*mvc*_: *β* = 4.51 [3.62, 5.40]) compared to higher torque levels (0.2 to 0.25 *τ*_*mvc*_: *β* = 2.94 [2.13, 3.74]).

**Figure 3:**
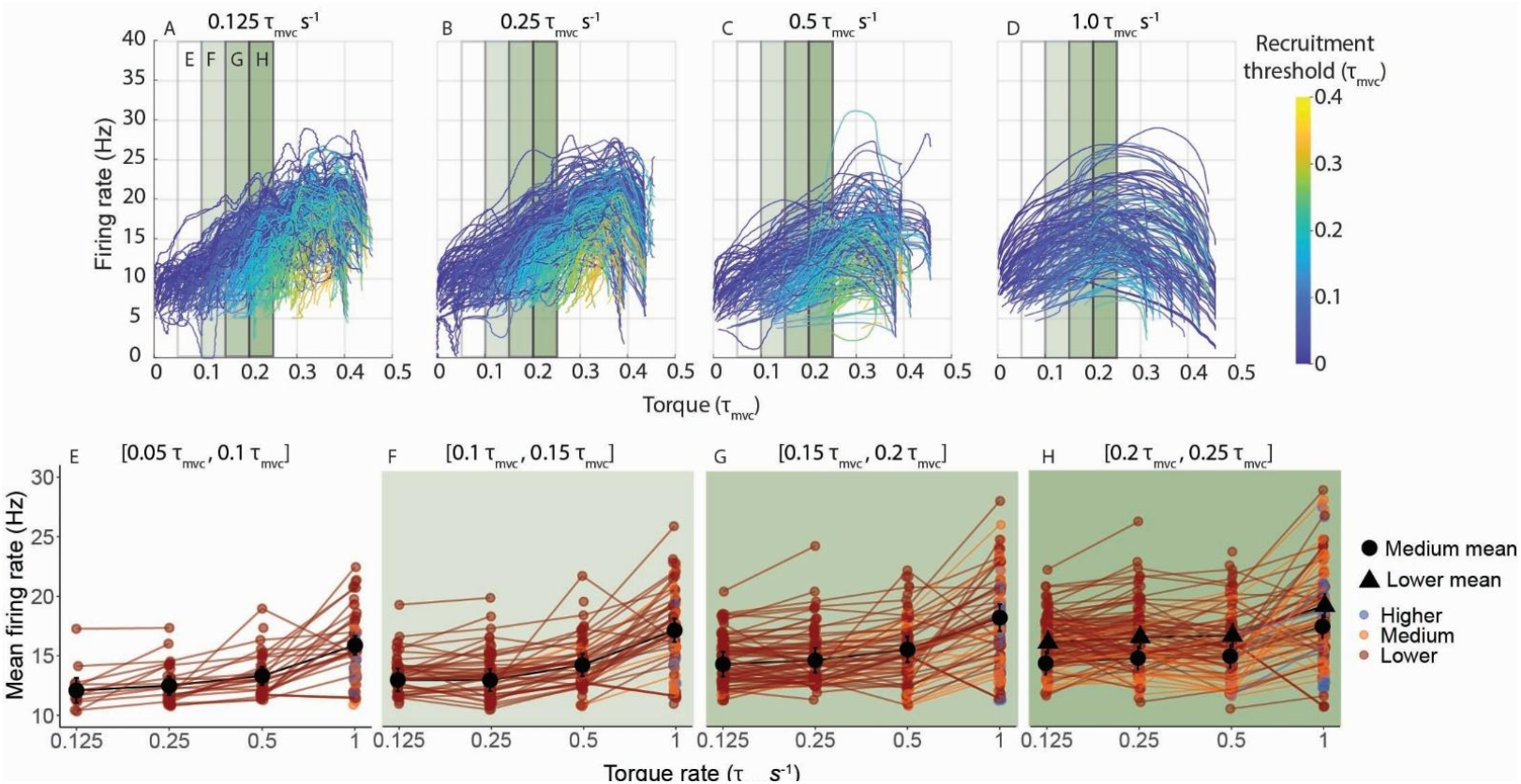
Mean motor unit firing rates increased with rate of torque development. Smoothed motor unit firing rates (n=263) are depicted for 0.125, 0.25, 0.5, and 1.0 τ_mvc_ s^−1^ conditions (A, B, C, D, respectively). From these data, the mean firing rates were taken in bins from 0.05 to 0.1 τ_mvc_, 0.1 to 0.15 τ_mvc_, 0.15 to 0.2 τ_mvc_, and 0.2 to 0.25 τ_mvc_. The green shaded regions, which represent different torque levels in A-D, corresponded to the means in E, F, G, H for all participants. Mean values with 95% confidence intervals are shown for the low threshold units (motor units recruited from 0 to 0.13 τ_mvc_). Lines in E-H represent tracked motor units (not all motor units were tracked across all conditions). Medium threshold units (0.13 to 0.27 τ_mvc_) are only reported in the 0.2 to 0.25 τ_mvc_ range given the low number detected at lower torque levels. Means are reported as estimated marginal means.

### 4.4 Fascicle dynamics

TA fascicles shortened by similar amounts but at greater shortening velocities with increases in rate of torque development. There was no significant difference in minimum fascicle strain with changes in contraction rate (*p* = 0.715, Figure 4A), with mean a peak fascicle strain of −0.14 [−0.16, −0.11] across all trials. Peak fascicle strain rate decreased (i.e., higher shortening rates) with rate of torque development (*p <* 0.001, Figure 4B). Pairwise comparisons showed the magnitude of peak fascicle strain rate increased (≈100%) from 0.31 [0.24, 0.37] *s*^−1^ to 0.63 [0.46, 0.79] *s*^−1^ (*p <* 0.001) from 0.125 *τ*_*mvc*_ *s*^−1^to the 1.0 *τ*_*mvc*_ *s*^−1^conditions. The relative timing of peak strain rates reported in Table 1 varied with rate of torque development with peaks occurring at torques of 0.06 [0.04, 0.09] *τ*_*mvc*_, 0.08 [0.05, 0.11] *τ*_*mvc*_, 0.11 [0.05, 0.16] *τ*_*mvc*_, and 0.16 [0.12, 0.21] *τ*_*mvc*_ for the 0.125, 0.25, 0.5, and 1.0 *τ*_*mvc*_ *s*^−1^conditions, respectively.

**Figure 4:**
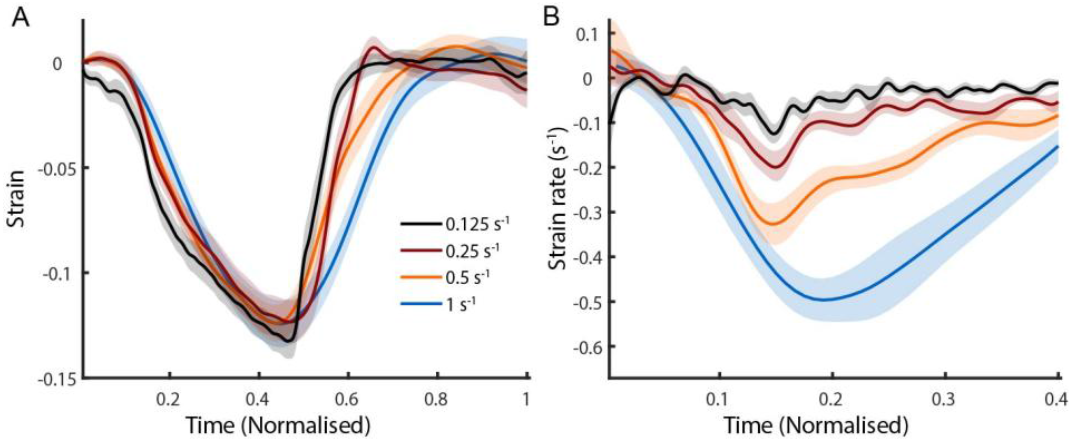
Tibialis anterior fascicles shortened by similar amounts but at higher strain rates as rate of torque development increased. Fascicle strain (A) and strain rates (B) vs. time normalised to cycle length. Shaded regions denote ± SE. Strain rates are shown only for the ramp period of the contraction and negative values indicate shortening.

### 4.5 Energetic rates

Energetic rates increased with rate of torque development (*p <* 0.001, Figure 5). The mean energetic rate was ≈220% higher in the fastest 1.0 *τ*_*mvc*_ *s*^−1^ condition (0.435 [0.321, 0.549] *W kg*^−1^) compared to the slowest 0.125 *τ*_*mvc*_ *s*^−1^condition (0.139 [0.022, 0.256] *W kg*^−1^; *p <* 0.001). Pairwise comparisons also revealed significant differences between the 0.125 *τ*_*mvc*_ *s*^−1^and the 0.5 *τ*_*mvc*_ *s*^−1^ condition (0.352 [0.237, 0.466] *W kg*^−1^; *p* = 0.005), and between the 0.25 *τ*_*mvc*_ *s*^−1^(0.0563 [0.094, 0.322] *W kg*^−1^) and the 1.0 *τ*_*mvc*_ *s*^−1^ condition (*p* = 0.002). No significant difference was observed between the two fastest torque rate conditions (0.5 *τ*_*mvc*_ *s*^−1^ and 1.0 *τ*_*mvc*_ *s*^−1^; *p* = 0.5).

**Figure 5:**
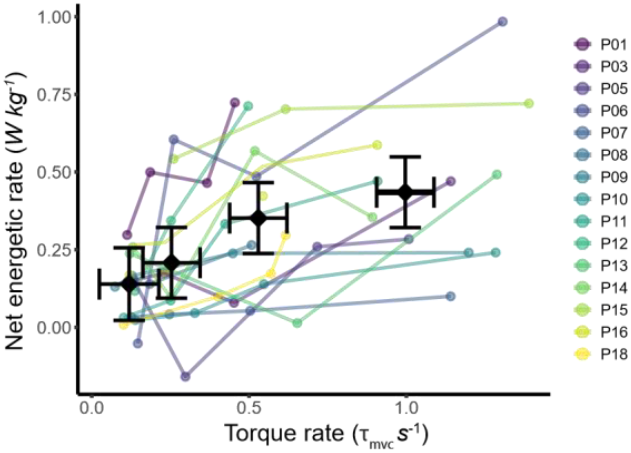
Energetic rates increased with rate of torque development. The mean (± 95% CIs) energetic rates are shown as a function of rate of torque development. Means are grouped based on condition (0.125, 0.25, 0.5, 1.0 τ_mvc_ s^−1^ ideal torque rates). Colours represent different participants. Means are reported as estimated marginal means.

## 5 Discussion

This study demonstrates changes in rate of torque development are associated not only with changes to muscle mechanical state, but also with changes in the neural control and energetic rates of muscle contraction. Simultaneous measures of *in vivo* motor unit discharge characteristics and muscle energy use provide insights into the relationship between neural control and energetics. The results support the hypothesis that higher rates of torque development are associated with increases in peak muscle strain rates, require higher neural drive and are associated with greater energetic rates. These results help to bridge the gap between idealised energy predictions based on isolated muscle preparations and real world *in vivo* muscle energetics [29], providing critical data for refining and validating mathematical models of skeletal muscle energy use.

### 5.1 Motor unit discharge characteristics

Decreases in recruitment threshold with increases in rate of torque development are consistent with previous studies [30,31], and are largely explained by the presence of series elasticity, including connective tissue and tendon. This elasticity allows the muscle fascicles to shorten despite fixed joint angles. Thus, to produce force at higher rates of torque development, muscle fibres must shorten at higher strain rates. Given the force-velocity constraints of skeletal muscle [32], higher neural drive is required to maintain a given force, leading to the recruitment of motor units at lower levels of torque with higher firing rates. Here, the differences in recruitment threshold with increasing rates of torque development were observed in motor units tracked across different conditions, ensuring that these differences reflect changes in the timing of the recruitment relative to joint torque, rather than the recruitment of different motor units with potentially different intrinsic properties. This supports the interpretation that a change in recruitment threshold is the consequence of higher neural drive at similar magnitudes of joint torque.

Previous studies have captured the role of rate of torque development on motor unit discharge characteristics, but have been limited in the number of motor units identified through intramuscular recordings [30,33] or have not captured muscle mechanical state over the duration of the contraction [31,34]. Our study demonstrates, through measuring large populations of motor units, that changes in motor unit discharge characteristics depend on the recruitment threshold, used here as a proxy of motor unit size. Specifically, recruitment thresholds of motor units recruited later during the contractions were more affected by the rate of force development (Figure 2). Analysis further revealed that the changes in mechanical state align with changes in motor unit rate coding. Specifically, with faster rates of torque development, peak muscle strain rates occur earlier in the contraction (during the ramp phase), which explains the higher level of neural drive that also occurs earlier in the ramp phase of the contraction cycle (both during the ramp phase and at recruitment (Figure 2C)). For instance, the peak strain rate in the 0.5 and 1.0 *τ*_*mvc*_ *s*^−1^ occurred at 0.11 [0.05, 0.16] *τ*_*mvc*_ and 0.16 [0.12, 0.21] *τ*_*mvc*_, which is within the ranges 0.1 to 0.15 *τ*_*mvc*_ and 0.15 to 0.2 *τ*_*mvc*_ where the largest fixed effects of rate of torque development on firing rates were detected (Figure 3FG). These higher motor unit firing rates observed early in the ramp phase at faster rates of force development align with previous findings demonstrating that the tibialis anterior relies mainly on rate coding to modulate force [21]. Of note, other lower limb muscles rely more on recruitment to modulate force (e.g. vastus lateralis [21]), which, together with muscle-dependent series elasticity, would likely lead to different energetic rates.

### 5.2 Muscle energetic rate

Higher energetic rates were observed with increases in rate of torque development, consistent with existing studies during knee extension tasks [12]. This study captures a range of mechanical demands on muscle energy use and neural control, including slow contractions (reflecting slow, controlled movements) to medium rates of contraction (reflecting walking) to faster contractions (reflecting running or ballistic movements) [35]. Across conditions, concurrent changes in both mechanical behaviour and neural control highlight their combined influence on muscle energy consumption.

Muscle mechanical state has been identified to be a prime determinant of muscle energy use through both *ex vivo* [2] and *in vivo* [14,36] experimental studies as well as with mathematical models [11,16]. The mechanics of muscle contraction can explain a substantial portion of the changes in energetic rates with rate of torque development as the energetic rate of cross-bridge cycling increases with faster fascicle strain rates [10,36]. Additionally, faster rates of torque development required more units to be active at lower torque levels (reduced recruitment threshold) at higher firing rates (Figure 2), which will also increase the energetic costs associated with muscle activation (primarily Ca^+2^ transport costs [37]).

Simultaneous measurement of muscle motor unit discharge characteristics and energetic rates provides an avenue to understand how *in vivo* recruitment patterns could influence muscle energy use [29]. Capturing recruitment strategies can be particularly important in dynamic contractions with fast muscle shortening rates. While we used low-to moderate intensity contractions to limit fatigue and ensure primarily aerobic pathways, we demonstrate earlier recruitment of higher threshold units with faster rates of torque development is required. This has consequences for muscle energy use as higher threshold motor units are associated with faster-type fibres [7]; if the contraction is at a sufficient intensity to recruit truly high threshold motor units with faster-type fibres, this could substantially influence muscle energetic rates. When it is possible to dissociate drive to low and high threshold motor units, changes in recruitment order may suggest a mechanism to improve contraction efficiency, as fast-type fibres can produce more force at higher strain rates [38], but this requires further exploration.

Motor unit discharge characteristics have been shown to vary with ageing and neurological disorders [39] potentially explaining some of the age- and disease-associated increases in movement costs [40]. The recruitment of different muscle fibre-types, as captured in existing mathematical models, can substantially influence energetic rates [8]. This study provides evidence that variations in mechanical demands lead to altered recruitment strategies with variable recruitment timing and discharge rates (Figures 2,3). The firing rate of a motor unit and the time active could influence the energetics associated with both Ca^+2^ dynamics [37] and the recovery time periods [41].

### 5.3 Inter-individual variability

Energetic rate measures, normalised to participant body mass, varied substantially across the healthy, similarly aged participants in this study (Figure 5). This is consistent with existing literature when individual data is reported (e.g. [13]). One possible explanation could be the variability in fascicle dynamics between participants with a coefficient of variation of 0.4 (Mean±SD: 0.130 ± 0.052, Table 1) in peak fascicle strain for the 1.0 *τ*_*mvc*_ *s*^−1^ condition; this is likely a result of differences in tendon compliance and muscle architecture, which has been shown to influence muscle energetics [42]. Differences in tibialis anterior muscle fibre-type could also play a role with substantial differences in efficiency from slower to faster fibres [6], which has been demonstrated to influence inter-individual variability in energetic rates [43].

### 5.4 Considerations and future directions

Indirect calorimetry has been used previously to measure individual muscle energetic rates (e.g. [12–14]), but as a whole-body measure, obtaining muscle-level estimates of energy use remains difficult. Due to the relative size of the tibialis anterior muscle, the muscle energetic rates were close to the resting rates and thus for some participants where obtaining a resting rate was difficult, the net rates were negative. Another difficulty in isolating the energetic cost from the tibialis anterior, is that it may only account for about 50% of the total ankle torque [44], making it possible that activation of other dorsiflexor muscles also contributed to the measured energetic rate. To better obtain muscle-specific energy estimates, other techniques such as functional phosphorus magnetic resonance spectroscopy [45,46] may provide more accurate measures, but concurrent recordings of motor unit activity within an MRI remains challenging.

Co-activation is a possible contributor to measured energetic rates; however, magnitudes of co-activation were low (Supplementary Figure S1) and it is likely EMG amplitude levels were overestimated due to normalisation (e.g. no twitch interpolation of the maximum voluntary contraction [47]) and crosstalk in the plantarflexor muscles [48]. Further, contributions to whole-body energetic rate were likely minimal as peak co-activation magnitude occurred in the fastest rate of torque development conditions, which had the briefest periods of co-activation. The torque integral is a potential contributor to the energetic rates due to the muscle relaxation time occupying a larger fraction of the contraction cycle at faster rates of torque development. While both co-activation and torque integral might increase energetic cost, in the fastest conditions (0.5 and 1.0 *τ*_*mvc*_ *s*^−1^) there was not a substantial increase in energy use compared to the lower rate of torque development. Less variation across the highest conditions could be due to difficulties in accurately measuring flow rates, or it is possible the greater time to relax the muscle reduced aerobic recovery time period within the contraction cycle.

Future work is needed to explicitly relate motor unit behaviour to muscle energetic rate, which likely warrants the development of a mathematical model accounting for both the nonlinear muscle mechanics and detailed motor unit level muscle control. The motor unit data obtained here can be used to inform motor unit pool models [49], while the fascicle dynamics can be used to inform models of muscle mechanics [50] which together characterise the mechanisms governing energy use. The energetics data can then be used to validate the resulting energetic predictions [16]. The modelling framework could isolate the relative contributions of neural control and mechanics on muscle energy use, while potentially capturing individual differences (e.g. fibre-type properties or individual fascicle dynamics) on muscle energy use.

## 6 Conclusion

This study simultaneously captured changes in muscle energy use and neural control strategies by varying the rate of torque development. We conclude that motor units are recruited earlier in the contraction with higher motor unit firing rates accompanying increases in fascicle strain rates. Altered motor unit discharge characteristics were concurrently measured alongside increases in muscle energy use highlighting the potential role of neural control strategies on muscle energetics. These data are critical for informing mathematical models to establish causal links between neural control, muscle mechanics, and energy use that can more accurately predict the energetic costs of movement.

## 7 Acknowledgements

The authors thank Gary Wilson for his assistance and expertise in setup of the indirect calorimetry system.

## References

1. McAllister MJ, Chen A, Selinger JC. 2025 Behavioural energetics in human locomotion: how energy use influences how we move. J. Exp. Biol. 228, JEB248125. (doi:10.1242/jeb.248125)

2. Woledge RC, Curtin NA, Homsher E. 1985 Energetic aspects of muscle contraction. Monogr. Physiol. Soc. 41, 1–357. (doi:10.1152/jappl.1986.60.3.1098)

3. Barclay CJ, Curtin NA. 2023 Advances in understanding the energetics of muscle contraction. J Biomech 156, 111669. (doi:10.1016/j.jbiomech.2023.111669)

4. Henneman E. 1957 Relation between size of neurons and their susceptibility to discharge. Science 126, 1345–1347. (doi:10.1126/SCIENCE.126.3287.1345)

5. Rattay F. 1990 Electrical nerve stimulation. Springer.

6. Barclay CJ. 2019 Efficiency of Skeletal Muscle. In Muscle and Exercise Physiology, pp. 111–127. Academic Press. (doi:10.1016/B978-0-12-814593-7.00006-2)

7. McPhedran AM, Wuerker RB, Henneman E. 1965 PROPERTIES OF MOTOR UNITS IN A HOMOGENEOUS RED MUSCLE (SOLEUS) OF THE CAT. J Neurophysiol 28, 71–84. (doi:10.1152/jn.1965.28.1.71)

8. Umberger BR, Gerritsen KGM, Martin PE. 2003 A Model of Human Muscle Energy Expenditure. Comp Meth Biomech Biomed Eng 6, 99–111. (doi:10.1080/1025584031000091678)

9. Bigland B, Lippold OCJ. 1954 The relation between force, velocity and integrated electrical activity in human muscles. J Physiol 123, 214–224. (doi:10.1113/jphysiol.1954.sp005044)

10. Fenn WO. 1923 A quantitative comparison between the energy liberated and the work performed by the isolated sartorius muscle of the frog. J Physiol 58, 175–203. (doi:10.1113/jphysiol.1923.sp002115)

11. Lichtwark GA, Wilson AM. 2005 A modified Hill muscle model that predicts muscle power output and efficiency during sinusoidal length changes. J Exp Biol 208, 2831–2843. (doi:10.1242/jeb.01709)

12. Van Der Zee TJ, Kuo AD. 2021 The high energetic cost of rapid force development in muscle. J. Exp. Biol. 224. (doi:10.1242/JEB.233965/237823)

13. Beck ON, Gosyne J, Franz JR, Sawicki GS. 2020 Cyclically producing the same average muscle-tendon force with a smaller duty increases metabolic rate. Proc R Soc B 287, 20200431. (doi:10.1098/rspb.2020.0431)

14. Beck ON, Trejo LH, Schroeder JN, Franz JR, Sawicki GS. 2022 Shorter muscle fascicle operating lengths increase the metabolic cost of cyclic force production. J Appl Physiol 133, 524–533. (doi:10.1152/japplphysiol.00720.2021)

15. Lentz-Nielsen N, Boysen MD, Munk-Hansen M, Laursen AD, Steffensen M, Engelund BK, Iversen K, Larsen RG, de Zee M. 2023 Validation of Metabolic Models for Estimation of Energy Expenditure During Isolated Concentric and Eccentric Muscle Contractions. J. Biomech. Eng. 145. (doi:10.1115/1.4063640)

16. Konno RN, Lichtwark GA, Dick TJM. 2025 Using physiologically based models to predict in vivo skeletal muscle energetics. J Exp Biol 228, jeb249966. (doi:10.1242/jeb.249966)

17. Callahan DM, Umberger BR, Kent-Braun JA. 2013 A Computational Model of Torque Generation: Neural, Contractile, Metabolic and Musculoskeletal Components. PLOS ONE 8, e56013. (doi:10.1371/JOURNAL.PONE.0056013)

18. Martinez-Valdes E et al. 2023 Consensus for experimental design in electromyography (CEDE) project: Single motor unit matrix. J Electromyogr Kinesiol 68, 102726. (doi:10.1016/j.jelekin.2022.102726)

19. Brand RA, Pedersen DR, Friederich JA. 1986 The sensitivity of muscle force predictions to changes in physiologic crosssectional area. J Biomech 19, 589–596. (doi:10.1016/0021-9290(86)90164-8)

20. De Zee M, Voigt M. 2002 Assessment of functional series elastic stiffness of human dorsiflexors with fast controlled releases. J Appl Physiol 93, 324–329. (doi:10.1152/japplphysiol.00696.2001)

21. Avrillon S, Hug F, Enoka RM, Caillet AH, Farina D. 2024 The identification of extensive samples of motor units in human muscles reveals diverse effects of neuromodulatory inputs on the rate coding. eLife 13, RP97085. (doi:10.7554/eLife.97085)

22. Avrillon S, Hug F, Baker SN, Gibbs C, Farina D. 2024 Tutorial on MUedit: An open-source software for identifying and analysing the discharge timing of motor units from electromyographic signals. J Electromyogr Kinesiol 77, 102886. (doi:10.1016/j.jelekin.2024.102886)

23. Del Vecchio A, Holobar A, Falla D, Felici F, Enoka RM, Farina D. 2020 Tutorial: Analysis of motor unit discharge characteristics from high-density surface EMG signals. J Electromyogr Kinesiol 53, 102426. (doi:10.1016/j.jelekin.2020.102426)

24. De Luca CJ, LeFever RS, McCue MP, Xenakis AP. 1982 Behaviour of human motor units in different muscles during linearly varying contractions. Physiol 329, 113–128. (doi:10.1113/jphysiol.1982.sp014293)

25. Van Der Zee TJ, Tecchio P, Hahn D, Raiteri BJ. 2025 UltraTimTrack: a Kalman-filter-based algorithm to track muscle fascicles in ultrasound image sequences. PeerJ Comp Sci 11, e2636. (doi:10.7717/peerj-cs.2636)

26. Gill PK, Kipp S, Beck ON, Kram R. 2023 It is time to abandon single-value oxygen uptake energy equivalents. J Appl Physiol 134, 887–890. (doi:10.1152/japplphysiol.00353.2022)

27. Péronnet F, Massicotte D. 1991 Table of nonprotein respiratory quotient: an update. Can. J. Sport Sci. J. Can. Sci. Sport 16, 23–29.

28. Lenth, RV. 2025 emmeans: Estimated Marginal Means, aka Least-Squares Means.

29. Lichtwark GA, Jessup LN, Konno RN, Riveros-Matthey CD, Dick TJM. 2025 Integrating muscle energetics into biomechanical models to understand variance in the cost of movement. J Exp Biol 228, JEB248022. (doi:10.1242/jeb.248022)

30. Aeles J, Bellett M, Lichtwark GA, Cresswell AG. 2022 The effect of small changes in rate of force development on muscle fascicle velocity and motor unit discharge behaviour. Eur J Appl Physiol 122, 1035–1044. (doi:10.1007/s00421-022-04905-7)

31. Chen C, Liu X, Qiu F. 2025 Characterization of motor unit activities during isometric elbow flexion with different speeds. J. NeuroEngineering Rehabil. 22, 236. (doi:10.1186/s12984-025-01765-y)

32. Hill A. 1938 The Heat of Shortening and the Dynamic Constants of Muscle. Proc R Soc B 126, 136–195.

33. Desmedt JE, Godaux E. 1977 Fast motor units are not preferentially activated in rapid voluntary contractions in man. Nat. 1977 2675613 267, 717–719. (doi:10.1038/267717a0)

34. Del Vecchio A, Negro F, Falla D, Bazzucchi I, Farina D, Felici F. 2018 Higher muscle fiber conduction velocity and early rate of torque development in chronically strength-trained individuals. J. Appl. Physiol. 125, 1218–1226. (doi:10.1152/JAPPLPHYSIOL.00025.2018/ASSET/IMAGES/LARGE/ZDG0101827480004.JPEG)

35. Cavagna GA, Heglund NC, Taylor CR. 1977 Mechanical work in terrestrial locomotion: two basic mechanisms for minimizing energy expenditure. Am. J. Physiol. - Regul. Integr. Comp. Physiol. 2. (doi:10.1152/ajpregu.1977.233.5.r243)

36. Ortega JO, Lindstedt SL, Nelson FE, Jubrias SA, Kushmerick MJ, Conley KE. 2015 Muscle force, work and cost: a novel technique to revisit the Fenn Effect. J Exp Biol, jeb.114512. (doi:10.1242/jeb.114512)

37. Barclay CJ, Woledge RC, Curtin NA. 2007 Energy turnover for Ca2+ cycling in skeletal muscle. J Muscle Res Cell Motil 28, 259–274. (doi:10.1007/s10974-007-9116-7)

38. Lai AKM, Biewener AA, Wakeling JM. 2018 Metabolic cost underlies task-dependent variations in motor unit recruitment. J R Soc Interface 15, 20180541. (doi:10.1098/rsif.2018.0541)

39. Hepple RT, Rice CL. 2016 Innervation and neuromuscular control in ageing skeletal muscle. J Physiol 594, 1965–1978. (doi:10.1113/JP270561)

40. Boyer KA et al. 2023 Age-related changes in gait biomechanics and their impact on the metabolic cost of walking: Report from a National Institute on Aging workshop. Exp. Gerontol. 173, 112102. (doi:10.1016/j.exger.2023.112102)

41. Phillips SK, Takei M, Yamada K. 1993 The time course of phosphate metabolites and intracellular pH using 31P NMR compared to recovery heat in rat soleus muscle. J. Physiol. 460, 693–704. (doi:10.1113/jphysiol.1993.sp019494)

42. Lichtwark GA, Barclay CJ. 2010 The influence of tendon compliance on muscle power output and efficiency during cyclic contractions. J Exp Biol 213, 707–714. (doi:10.1242/jeb.038026)

43. Swinnen W, Lievens E, Hoogkamer W, De Groote F, Derave W, Vanwanseele B. 2024 Inter-Individual Variability in Muscle Fiber–Type Distribution Affects Running Economy but Not Running Gait at Submaximal Running Speeds. Scand. J. Med. Sci. Sports 34, e14748. (doi:10.1111/SMS.14748)

44. Raiteri BJ, Lauret L, Hahn D. 2023 The force-length relation of the young adult human tibialis anterior. PeerJ 11, e15693. (doi:10.7717/PEERJ.15693/SUPP-6)

45. Haeufle DFB, Siegel J, Hochstein S, Gussew A, Schmitt S, Siebert T, Rzanny R, Reichenbach JR, Stutzig N. 2020 Energy Expenditure of Dynamic Submaximal Human Plantarflexion Movements: Model Prediction and Validation by in-vivo Magnetic Resonance Spectroscopy. Front Bioeng Biotech 8. (doi:10.3389/FBIOE.2020.00622/FULL)

46. Boska M. 1994 ATP production rates as a function of force level in the human gastrocnemius/soleus using^31^ P MRS. Magn Reson Med 32, 1–10. (doi:10.1002/mrm.1910320102)

47. Belanger AY, McComas AJ. 1981 Extent of motor unit activation during effort. J Appl Physiol 51, 1131–1135. (doi:10.1152/jappl.1981.51.5.1131)

48. Raiteri BJ, Hug F, Cresswell AG, Lichtwark GA. 2016 Quantification of muscle co-contraction using supersonic shear wave imaging. J Biomech 49, 493–495. (doi:10.1016/j.jbiomech.2015.12.039)

49. Caillet AH, Phillips ATM, Farina D, Modenese L. 2022 Estimation of the firing behaviour of a complete motoneuron pool by combining electromyography signal decomposition and realistic motoneuron modelling. PLoS Comput Biol 18, e1010556. (doi:10.1371/journal.pcbi.1010556)

50. Zajac FE. 1989 Muscle and tendon: properties, models, scaling, and application to biomechanics and motor control. Crit. Rev. Biomed. Eng. 17, 359–411.

