## Supplementary Material for "The impact of mechanical requirements on the neural control of skeletal muscle and subsequent energetic rates"

### 1 SI: Co-activation

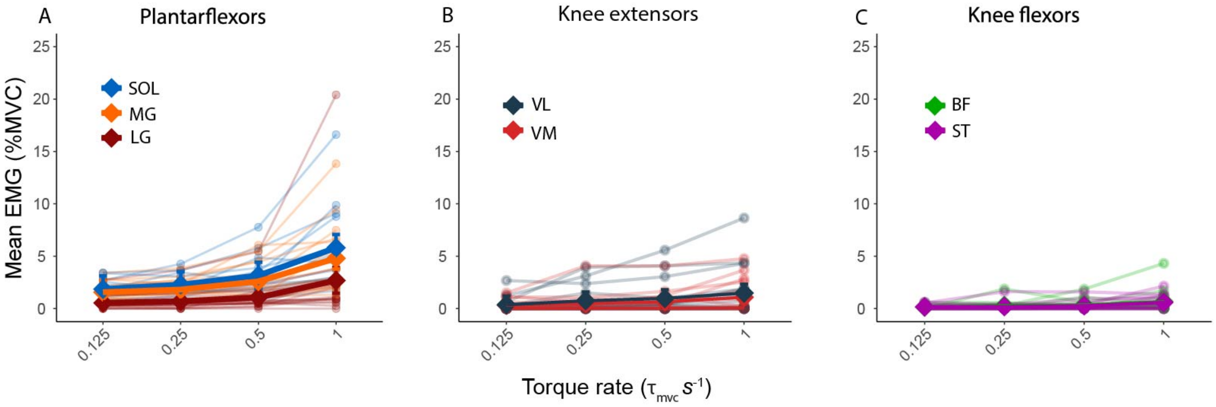


***Figure S1: Mean normalised global EMG across trials.*** *MG, SOL were computed by a differential using the monopolar signals from the HDsEMG grids, while LG, VL, VM, BF, ST were computed from the differential sEMG. Mean EMG of all muscles increased with rate of torque development ($p < 0.001$), but with low levels of co-activation ($< 5.7\%$ MVC in the plantarflextors, $< 0.02\%$ in the knee extensors and flexors). SOL: soleus, MG: medial gastronemius, LG: lateral gastronemius, VL: vastus lateralis, VM: vastus medialis, BF: biceps femoris, ST: semitendinosis. Colours represent different participants. Means were calculated as estimated marginal means with error bars representing 95% CI.*

### 2 SI: Mean values and post-hoc comparisons

#### 2.1 Torque integral comparisons

***Table S1: Pairwise comparisons of torque integral estimated marginal means across thresholds with Tukey adjustment.*** *Torque integral was computed as the integral of the torque over the whole trial. Contrasts represent the rate of torque development conditions compared, while estimates represent the difference in estimated marginal means. SE: standard error.*

| **Contrast** ($\tau_{mvc} s^{-1}$) | **Estimate** ($\tau_{mvc}\, s$) | **SE** ($\tau_{mvc}\, s$) | **t ratio** | **p value** |
|---|---:|---:|---:|---:|
| 0.125 → 0.25 | -3.69 | 1.07 | -3.451 | 0.006 |
| 0.125 → 0.5 | -9.74 | 1.07 | -9.106 | <0.001 |
| 0.125 → 1 | -17.59 | 1.07 | -16.443 | <0.001 |
| 0.25 → 0.5 | -6.05 | 1.07 | -5.655 | <0.001 |
| 0.25 → 1 | -13.89 | 1.07 | -12.992 | <0.001 |
| 0.5 → 1 | -7.85 | 1.07 | -7.337 | <0.001 |

#### 2.2 Global surface electromyography pairwise comparisons

***Table S2: Pairwise comparisons of maximum EMG amplitude estimated marginal means in the tibialis anterior across thresholds with Tukey adjustment.*** *Maximum EMG amplitude was computed as the maximum EMG amplitude over the contraction cycle. Contrasts represent the rate of torque development conditions compared, while estimates represent the difference in estimated marginal means. SE: standard error.*

| **Contrast** ($\tau_{mvc}s^{-1}$) | **Estimate** (Normalised) | **SE** (Normalised) | **t ratio** | **p value** |
|---|---:|---:|---:|---:|
| 0.125 → 0.25 | 0.00636 | 0.0295 | 0.215 | 0.996 |
| 0.125 → 0.5 | -0.01056 | 0.0295 | -0.358 | 0.984 |
| 0.125 → 1 | -0.13870 | 0.0295 | -4.697 | <0.001 |
| 0.25 → 0.5 | -0.01692 | 0.0295 | -0.573 | 0.940 |
| 0.25 → 1 | -0.14506 | 0.0295 | -4.912 | <0.001 |
| 0.5 → 1 | -0.12814 | 0.0295 | -4.339 | <0.001 |

***Table S3: Estimated marginal means for mean global EMG amplitudes with 95% confidence intervals across muscles muscles and rate of torque development.*** *Mean global EMG amplitude was computed as the mean value over the contraction cycle including both the rest period and the contraction. Values are reported as a percentage of the maximum EMG amplitude value in the MVC trial. SOL: soleus, MG: medial gastronemius, LG: lateral gastronemius, VL: vastus lateralis, VM: vastus medialis, BF: biceps femoris, ST: semitendinosis SE: standard error.*

| Muscle | Torque rate ($\tau_{mvc}s^{-1}$) | EMG Amplitude [95% CI] (%MVC) |
|---|---|---|
| VM | 0.125 | 0.325 [-0.240, 0.890] |
| VM | 0.25 | 0.489 [-0.076, 1.050] |
| VM | 0.5 | 0.644 [0.079, 1.210] |
| VM | 1 | 1.080 [0.512, 1.640] |
| VL | 0.125 | 0.362 [-0.485, 1.210] |
| VL | 0.25 | 0.749 [-0.098, 1.600] |
| VL | 0.5 | 0.967 [0.119, 1.810] |
| VL | 1 | 1.480 [0.631, 2.330] |
| BF | 0.125 | 0.152 [-0.192, 0.497] |
| BF | 0.25 | 0.286 [-0.058, 0.631] |
| BF | 0.5 | 0.350 [0.005, 0.695] |
| BF | 1 | 0.661 [0.316, 1.010] |
| ST | 0.125 | 0.151 [-0.062, 0.365] |
| ST | 0.25 | 0.191 [-0.022, 0.405] |
| ST | 0.5 | 0.272 [0.059, 0.486] |
| ST | 1 | 0.591 [0.377, 0.804] |
| LG | 0.125 | 0.538 [-0.685, 1.760] |
| LG | 0.25 | 0.691 [-0.531, 1.910] |
| LG | 0.5 | 1.060 [-0.160, 2.280] |
| LG | 1 | 2.680 [1.450, 3.900] |
| TA | 0.125 | 8.970 [6.870, 11.100] |
| TA | 0.25 | 10.400 [8.310, 12.500] |
| TA | 0.5 | 12.600 [10.400, 14.700] |
| TA | 1 | 17.400 [15.300, 19.500] |
| MG | 0.125 | 1.560 [0.540, 2.580] |
| MG | 0.25 | 1.810 [0.791, 2.830] |
| MG | 0.5 | 2.590 [1.570, 3.610] |
| MG | 1 | 4.800 [3.780, 5.820] |
| SOL | 0.125 | 1.850 [0.591, 3.120] |
| SOL | 0.25 | 2.280 [1.010, 3.540] |
| SOL | 0.5 | 3.150 [1.890, 4.420] |
| SOL | 1 | 5.810 [4.540, 7.070] |

***Table S4: Pairwise comparisons of global EMG amplitude estimated marginal means across thresholds with Tukey adjustment.*** *Mean global EMG amplitude was computed as the mean value over the contraction cycle including both the rest period and the contraction. Values are reported a percentage of the maximum EMG amplitude value in the MVC trial. Contrasts represent the rate of torque development conditions compared, while estimates represent the difference in estimated marginal means. SOL: soleus, MG: medial gastronemius, LG: lateral gastronemius, VL: vastus lateralis, VM: vastus medialis, BF: biceps femoris, ST: semitendinosis, SE: standard error.*

| Muscle | Contrast ($\tau_{mvc}s^{-1}$) | Estimate (%MVC) | SE (%MVC) | t ratio | p value |
|---|---|---|---|---|---|
| VM | 0.125 → 0.25 | -0.164 | 0.219 | -0.750 | 0.876 |
| VM | 0.125 → 0.5 | -0.319 | 0.219 | -1.460 | 0.470 |
| VM | 0.125 → 1 | -0.752 | 0.219 | -3.440 | 0.007 |
| VM | 0.25 → 0.5 | -0.155 | 0.219 | -0.709 | 0.893 |
| VM | 0.25 → 1 | -0.588 | 0.219 | -2.690 | 0.048 |
| VM | 0.5 → 1 | -0.433 | 0.219 | -1.980 | 0.211 |
| VL | 0.125 → 0.25 | -0.387 | 0.318 | -1.220 | 0.620 |
| VL | 0.125 → 0.5 | -0.604 | 0.318 | -1.900 | 0.242 |
| VL | 0.125 → 1 | -1.120 | 0.318 | -3.510 | 0.006 |
| VL | 0.25 → 0.5 | -0.218 | 0.318 | -0.684 | 0.903 |
| VL | 0.25 → 1 | -0.729 | 0.318 | -2.290 | 0.115 |
| VL | 0.5 → 1 | -0.512 | 0.318 | -1.610 | 0.384 |
| BF | 0.125 → 0.25 | -0.134 | 0.207 | -0.646 | 0.916 |
| BF | 0.125 → 0.5 | -0.198 | 0.207 | -0.954 | 0.776 |
| BF | 0.125 → 1 | -0.509 | 0.207 | -2.450 | 0.082 |
| BF | 0.25 → 0.5 | -0.064 | 0.207 | -0.308 | 0.990 |
| BF | 0.25 → 1 | -0.375 | 0.207 | -1.810 | 0.284 |
| BF | 0.5 → 1 | -0.311 | 0.207 | -1.500 | 0.446 |
| ST | 0.125 → 0.25 | -0.040 | 0.107 | -0.376 | 0.982 |
| ST | 0.125 → 0.5 | -0.121 | 0.107 | -1.130 | 0.670 |
| ST | 0.125 → 1 | -0.439 | 0.107 | -4.120 | 0.001 |
| ST | 0.25 → 0.5 | -0.081 | 0.107 | -0.759 | 0.872 |
| ST | 0.25 → 1 | -0.399 | 0.107 | -3.750 | 0.003 |
| ST | 0.5 → 1 | -0.318 | 0.107 | -2.990 | 0.022 |
| LG | 0.125 → 0.25 | -0.153 | 0.666 | -0.230 | 0.996 |
| LG | 0.125 → 0.5 | -0.525 | 0.666 | -0.788 | 0.859 |
| LG | 0.125 → 1 | -2.140 | 0.666 | -3.210 | 0.012 |
| LG | 0.25 → 0.5 | -0.372 | 0.666 | -0.558 | 0.944 |
| LG | 0.25 → 1 | -1.990 | 0.666 | -2.980 | 0.023 |
| LG | 0.5 → 1 | -1.610 | 0.666 | -2.420 | 0.086 |
| TA | 0.125 → 0.25 | -1.440 | 0.880 | -1.640 | 0.368 |
| TA | 0.125 → 0.5 | -3.580 | 0.880 | -4.070 | 0.001 |
| TA | 0.125 → 1 | -8.410 | 0.880 | -9.550 | <0.001 |
| TA | 0.25 → 0.5 | -2.140 | 0.880 | -2.430 | 0.085 |
| TA | 0.25 → 1 | -6.970 | 0.880 | -7.910 | <0.001 |
| TA | 0.5 → 1 | -4.830 | 0.880 | -5.480 | <0.001 |
| MG | 0.125 → 0.25 | -0.251 | 0.477 | -0.527 | 0.952 |
| MG | 0.125 → 0.5 | -1.030 | 0.477 | -2.170 | 0.149 |
| MG | 0.125 → 1 | -3.240 | 0.477 | -6.780 | <0.001 |
| MG | 0.25 → 0.5 | -0.782 | 0.477 | -1.640 | 0.368 |
| MG | 0.25 → 1 | -2.990 | 0.477 | -6.260 | <0.001 |
| MG | 0.5 → 1 | -2.200 | 0.477 | -4.620 | <0.001 |
| SOL | 0.125 → 0.25 | -0.422 | 0.651 | -0.648 | 0.916 |
| SOL | 0.125 → 0.5 | -1.300 | 0.651 | -1.990 | 0.208 |
| SOL | 0.125 → 1 | -3.950 | 0.651 | -6.070 | <0.001 |
| SOL | 0.25 → 0.5 | -0.876 | 0.651 | -1.340 | 0.541 |
| SOL | 0.25 → 1 | -3.530 | 0.651 | -5.420 | <0.001 |
| SOL | 0.5 → 1 | -2.650 | 0.651 | -4.080 | 0.001 |

#### 2.3 Motor unit discharge characteristics means and pairwise comparisons

##### 2.3.1 Recruitment threshold comparisons

***Table S5: Motor unit recruitment threshold estimated marginal means with 95% confidence intervals across different thresholds.*** *Recruitment thresholds were calculated as the torque at first motor unit discharge. The motor unit threshold category was determined based on recruitment threshold with lower being 0 to 0.13* $\tau_{mvc}$*, medium being 0.13 to 0.27* $\tau_{mvc}$*, and higher being 0.27 to 0.4* $\tau_{mvc}$*. Values were normalised to MVC torque.*

| **Torque rate** ($\tau_{mvc}s^{-1}$) | **Recruitment threshold [95% CI]** ($\tau_{mvc}$) |
|---|---|
| **Threshold category = Higher** | |
| 0.125 | 0.31 [0.29, 0.33] |
| 0.25 | 0.27 [0.26, 0.29] |
| 0.5 | 0.24 [0.23, 0.26] |
| 1 | 0.11 [0.09, 0.13] |
| **Threshold category = Medium** | |
| 0.125 | 0.20 [0.19, 0.22] |
| 0.25 | 0.17 [0.16, 0.19] |
| 0.5 | 0.15 [0.13, 0.16] |
| 1 | 0.09 [0.07, 0.10] |
| **Threshold category = Lower** | |
| 0.125 | 0.06 [0.05, 0.08] |
| 0.25 | 0.05 [0.04, 0.06] |
| 0.5 | 0.04 [0.03, 0.06] |
| 1 | 0.04 [0.02, 0.05] |

***Table S6: Pairwise comparisons of recruitment threshold estimated marginal means across rates with Tukey adjustment.*** *Contrasts are performed with in threshold categories and represent the rate of torque development conditions compared, while estimates represent the difference in estimated marginal means. SE: standard error.*

| **Threshold category** | **Contrast** ($\tau_{mvc}s^{-1}$) | **Estimate** ($\tau_{mvc}$) | **SE** ($\tau_{mvc}$) | **t ratio** | **p value** |
|---|---|---|---|---|---|
| Higher | $0.125 \rightarrow 0.25$ | 0.037 | 0.012 | 3.105 | 0.011 |
| Higher | $0.125 \rightarrow 0.5$ | 0.068 | 0.012 | 5.775 | <0.001 |
| Higher | $0.125 \rightarrow 1$ | 0.198 | 0.012 | 16.518 | <0.001 |
| Higher | $0.25 \rightarrow 0.5$ | 0.031 | 0.011 | 2.818 | 0.026 |
| Higher | $0.25 \rightarrow 1$ | 0.162 | 0.011 | 14.231 | <0.001 |
| Higher | $0.5 \rightarrow 1$ | 0.130 | 0.011 | 11.482 | <0.001 |
| Medium | $0.125 \rightarrow 0.25$ | 0.032 | 0.008 | 4.124 | <0.001 |
| Medium | $0.125 \rightarrow 0.5$ | 0.057 | 0.008 | 7.032 | <0.001 |
| Medium | $0.125 \rightarrow 1$ | 0.119 | 0.009 | 13.393 | <0.001 |
| Medium | $0.25 \rightarrow 0.5$ | 0.025 | 0.008 | 3.082 | 0.011 |
| Medium | $0.25 \rightarrow 1$ | 0.087 | 0.009 | 9.771 | <0.001 |
| Medium | $0.5 \rightarrow 1$ | 0.062 | 0.009 | 6.796 | <0.001 |
| Lower | $0.125 \rightarrow 0.25$ | 0.012 | 0.007 | 1.841 | 0.255 |
| Lower | $0.125 \rightarrow 0.5$ | 0.020 | 0.007 | 2.874 | 0.022 |
| Lower | $0.125 \rightarrow 1$ | 0.023 | 0.008 | 2.980 | 0.016 |
| Lower | $0.25 \rightarrow 0.5$ | 0.008 | 0.007 | 1.181 | 0.639 |
| Lower | $0.25 \rightarrow 1$ | 0.011 | 0.008 | 1.433 | 0.479 |
| Lower | $0.5 \rightarrow 1$ | 0.003 | 0.008 | 0.337 | 0.987 |

***Table S7: Pairwise comparisons of recruitment threshold estimated marginal means across threshold category with Tukey adjustment.*** *Contrasts are performed within torque rate conditions and represent the threshold categories compared, while estimates represent the difference in estimated marginal means. SE: standard error.*

| **Torque rate** ($\tau_{mvc}s^{-1}$) | **Contrast** | **Estimate** ($\tau_{mvc}$) | **SE** ($\tau_{mvc}$) | **t ratio** | **p value** |
|---|---|---|---|---|---|
| 0.125 | Higher → Medium | 0.107 | 0.010 | 10.253 | <0.001 |
| 0.125 | Higher → Lower | 0.249 | 0.010 | 24.806 | <0.001 |
| 0.125 | Medium → Lower | 0.142 | 0.007 | 19.323 | <0.001 |
| 0.25 | Higher → Medium | 0.102 | 0.010 | 10.574 | <0.001 |
| 0.25 | Higher → Lower | 0.224 | 0.009 | 24.147 | <0.001 |
| 0.25 | Medium → Lower | 0.122 | 0.007 | 16.644 | <0.001 |
| 0.5 | Higher → Medium | 0.096 | 0.010 | 9.635 | <0.001 |
| 0.5 | Higher → Lower | 0.201 | 0.010 | 20.726 | <0.001 |
| 0.5 | Medium → Lower | 0.105 | 0.008 | 12.868 | <0.001 |
| 1 | Higher → Medium | 0.028 | 0.011 | 2.553 | 0.029 |
| 1 | Higher → Lower | 0.074 | 0.010 | 7.035 | <0.001 |
| 1 | Medium → Lower | 0.046 | 0.009 | 4.846 | <0.001 |

###### 2.3.2 Firing rate at recruitment

***Table S8: Estimated marginal means of firing rates at recruitment with 95% confidence intervals.*** *Firing rates at recruitment are calculated as the mean firing rate over the first 4 instantaneous motor unit discharges.*

| **Torque rate** ($\tau_{mvc}s^{-1}$) | **Firing rate [95% CI]** ($Hz$) |
|---|---|
| **Threshold category = Higher** | |
| 0.125 | 12.8 [11.48, 14.10] |
| 0.25 | 14.4 [13.17, 15.60] |
| 0.5 | 14.5 [13.34, 15.70] |
| 1 | 18.9 [17.69, 20.20] |
| **Threshold category = Medium** | |
| 0.125 | 11.9 [10.85, 12.90] |
| 0.25 | 13.1 [12.08, 14.10] |
| 0.5 | 14.0 [12.93, 15.00] |
| 1 | 17.5 [16.38, 18.60] |
| **Threshold category = Lower** | |
| 0.125 | 10.3 [9.35, 11.30] |
| 0.25 | 11.7 [10.74, 12.60] |
| 0.5 | 14.1 [13.07, 15.10] |
| 1 | 19.4 [18.32, 20.50] |

***Table S9: Pairwise comparisons of firing rate at recruitment within each threshold level.*** *Contrasts are performed with in threshold categories and represent the rate of torque development conditions compared, while estimates represent the difference in estimated marginal means. SE: standard error.*

| **Threshold category** | **Contrast** ($\tau_{mvc}s^{-1}$) | **Estimate** (Hz) | **SE** (Hz) | **t ratio** | **p value** |
|---|---|---|---|---|---|
| Higher | 0.125 → 0.25 | -1.62 | 0.72 | -2.238 | 0.114 |
| Higher | 0.125 → 0.5 | -1.78 | 0.72 | -2.466 | 0.066 |
| Higher | 0.125 → 1 | -6.16 | 0.74 | -8.363 | <0.001 |
| Higher | 0.25 → 0.5 | -0.16 | 0.68 | -0.236 | 0.995 |
| Higher | 0.25 → 1 | -4.54 | 0.70 | -6.517 | <0.001 |
| Higher | 0.5 → 1 | -4.38 | 0.70 | -6.289 | <0.001 |
| Medium | 0.125 → 0.25 | -1.23 | 0.48 | -2.575 | 0.050 |
| Medium | 0.125 → 0.5 | -2.12 | 0.50 | -4.244 | <0.001 |
| Medium | 0.125 → 1 | -5.65 | 0.55 | -10.353 | <0.001 |
| Medium | 0.25 → 0.5 | -0.88 | 0.50 | -1.778 | 0.285 |
| Medium | 0.25 → 1 | -4.41 | 0.55 | -8.094 | <0.001 |
| Medium | 0.5 → 1 | -3.53 | 0.56 | -6.322 | <0.001 |
| Lower | 0.125 → 0.25 | -1.39 | 0.40 | -3.490 | 0.003 |
| Lower | 0.125 → 0.5 | -3.78 | 0.43 | -8.743 | <0.001 |
| Lower | 0.125 → 1 | -9.10 | 0.47 | -19.220 | <0.001 |
| Lower | 0.25 → 0.5 | -2.39 | 0.43 | -5.546 | <0.001 |
| Lower | 0.25 → 1 | -7.70 | 0.47 | -16.347 | <0.001 |
| Lower | 0.5 → 1 | -5.31 | 0.49 | -10.751 | <0.001 |

***Table S10: Pairwise comparisons of firing rates at recruitment within each rate of torque development level.*** *Contrasts are performed within torque rate conditions and represent the threshold categories compared, while estimates represent the difference in estimated marginal means. SE: standard error.*

| **Rates** ($\tau_{mvc}s^{-1}$) | **Contrast** | **Estimate** (Hz) | **SE** (Hz) | **t ratio** | **p value** |
|---|---|---|---|---|---|
| 0.125 | Higher → Medium | 0.92 | 0.64 | 1.433 | 0.324 |
| 0.125 | Higher → Lower | 2.47 | 0.62 | 4.008 | <0.001 |
| 0.125 | Medium → Lower | 1.55 | 0.45 | 3.438 | 0.002 |
| 0.25 | Higher → Medium | 1.30 | 0.60 | 2.184 | 0.075 |
| 0.25 | Higher → Lower | 2.69 | 0.57 | 4.723 | <0.001 |
| 0.25 | Medium → Lower | 1.39 | 0.45 | 3.098 | 0.006 |
| 0.5 | Higher → Medium | 0.58 | 0.61 | 0.943 | 0.613 |
| 0.5 | Higher → Lower | 0.46 | 0.60 | 0.776 | 0.718 |
| 0.5 | Medium → Lower | -0.12 | 0.50 | -0.230 | 0.971 |
| 1 | Higher → Medium | 1.43 | 0.66 | 2.151 | 0.081 |
| 1 | Higher → Lower | -0.47 | 0.64 | -0.735 | 0.743 |
| 1 | Medium → Lower | -1.90 | 0.58 | -3.265 | 0.003 |

#### 2.4 Fascicle dynamics pairwise comparisons

***Table S11: Pairwise comparisons of peak fascicle strain rates with estimated marginal means across rates of torque development with Tukey adjustment.*** *Contrasts represent the rate of torque development conditions compared, while estimates represent the difference in estimated marginal means. Mean values are in the main text. SE: standard error.*

| **Contrast** | $(\tau_{mvc}s^{-1})$ | **Estimate** $(s^{-1})$ | **SE** $(s^{-1})$ | **t ratio** | **p value** |
|---|---|---|---|---|---|
| 0.125 → 0.25 | 0.01 | 0.07 | 42.2 | 0.086 | 0.999 |
| 0.125 → 0.5 | 0.10 | 0.07 | 42.2 | 1.546 | 0.420 |
| 0.125 → 1 | 0.32 | 0.07 | 42.2 | 4.784 | <0.001 |
| 0.25 → 0.5 | 0.10 | 0.07 | 42.2 | 1.460 | 0.470 |
| 0.25 → 1 | 0.31 | 0.07 | 42.2 | 4.698 | <0.001 |
| 0.5 → 1 | 0.22 | 0.07 | 42.2 | 3.238 | 0.012 |

#### 2.5 Energetic rate pairwise comparisons

***Table S12: Pairwise comparisons of net energetic rates across rates of torque development with Tukey adjustment.*** *Contrasts represent the rate of torque development conditions compared, while estimates represent the difference in estimated marginal means. Mean values are in the main text. SE: standard error.*

| **Contrast** ($\tau_{mvc} s^{-1}$) | **Estimate** ($W\,kg^{-1}$) | **SE** ($W\,kg^{-1}$) | **t ratio** | **p value** |
|---|---|---|---|---|
| 0.125 → 0.25 | -0.0686 | 0.0604 | -1.136 | 0.6693 |
| 0.125 → 0.5 | -0.2125 | 0.0597 | -3.562 | 0.0049 |
| 0.125 → 1 | -0.2956 | 0.0604 | -4.895 | 0.0001 |
| 0.25 → 0.5 | -0.1439 | 0.0589 | -2.441 | 0.0842 |
| 0.25 → 1 | -0.2270 | 0.0588 | -3.859 | 0.0020 |
| 0.5 → 1 | -0.0831 | 0.0589 | -1.411 | 0.4996 |
